# Allergen Extraction: Factors Influencing the Yield, Allergenicity and Sensitivity of Immunoassays

**DOI:** 10.1101/2020.02.20.957373

**Authors:** João Ricardo Almeida Soares, Isabelle Mazza Guimarães, Ana Luísa de Souza Oliveira, Claudia Regina Josetti das Neves Faccini, Erika Bertozzi de Aquino Mattos, Sónia Kristy Pinto Melo Rodrigues, Airton Pereira e Silva, Bárbara Oliveira Marmello, Gerlinde Agate Platais Brasil Teixeira

**Affiliations:** Gastrointestinal Immunology Group, Department of Immunobiology, Institute of Biology, Federal Fluminense University, Niterói, Rio de Janeiro 24020-150; Graduate Program in Science and Biotechnology, Federal Fluminense University, Niterói, Rio de Janeiro 24020-141, Brazil; Graduate Program in Pathology, Medicine School, Antônio Pedro Hospital, Federal Fluminense University, Niterói, Rio de Janeiro 24033-900

## Abstract

Food allergy is an IgE and or IgG immune-mediated reaction to food antigens. Knowledge of the allergenicity properties of proteins, how they react in the body and in diagnostic tests is necessary to adequately assess the allergenic potential of both natural foods and those produced through biotechnological processes. Thus, our aim was analyze the factors those influence the protein extraction of foods in terms of yield, allergenicity and sensitivity in immunoassays. Peanut proteins were extracted using 4 distinct extraction buffers (physiological saline, tris buffer, borate buffer with and without β-mercaptoethanol). The protein concentration of the obtained extracts was determined by the Lowry method. Polyacrylamide electrophoresis (SDS-PAGE) was used to compare the protein profile of each extract. Immunogenicity of each extract was verified by sensitizing two mouse strains (Balb/c and C57/BL6) with 100μg of the proteins, and the immunoreactivity was determined by ELISA. Our results show that extraction with the distinct buffers resulted in protein solutions with different yields and profiles. The antigenicity of the different extracts in a murine model of systemic immunization also demonstrated distinct patterns that varied depending on the extraction methods and mouse strain. The immunoreactivity varied in accordance to the protein extract used to coat the microtitration plates. In conclusion, **t**he protein yield and profile in the extracts is critically influenced by the salt composition and pH of the extraction buffers. This in turn influences both in vivo immunogenicity and in vitro immunoreactivity.

## Introduction

Considering that 4% to 8% of children present allergies and that a significant portion of these present symptoms that prevent school attendance which in turn takes, at least one of their caregivers away from work we can say that allergies have a high social impact [1]. In addition to absence at work, altering meal preparation routine and other social activities of the family negatively impacts stress levels [2]. Thus, adequate diagnosis is fundamental.

The best-known measure to prevent food allergy symptomatology in allergic people is the adoption of exclusion diets. To ensure clearer food labeling many countries have food allergen labeling laws. For example, the Food Allergen Labeling and Consumer Protection Act (FALCPA/USA-2004) requires that foods are labeled to identify the eight major food allergens: milk, egg, fish, crustacean-shellfish, tree nuts, wheat, peanuts and soybeans which account for over 90% of all documented food allergies in the U.S. [3]. Along with these 8 the Australian law requires the labeling of two other foods – sesame and lupin [4]. In the European Union 14 foods are on the required labeling list: the ten previously cited plus celery, mustard, Sulphur dioxide, and mollusks.

Regardless of whether ingested food is derived from animals or plants, the food matrix is composed of varying amounts of proteins, carbohydrates and fats. To perform reliable analysis of each macronutrient, several techniques have been developed to obtain adequate quantities of purified material with a minimum of structural and functional loss. For this, the main steps are extraction, quantification, biological function determination, sequence and structure identification [5]. Among the resources used in food protein analysis, the first step is the physical fragmentation of solid foods. This increases the contact surface of the food with the buffers during incubation, permitting a more efficient extraction [6]. Acidic, neutral or alkaline molecules are preferentially extracted when specific buffers with different salts and ionic strengths are employed. The efficiency of these buffers may vary according to the food matrix, thus altering the protein yield resulting from the extraction process [7]. Once a protein extract is obtained, quantification is a fundamental procedure influencing the registered protein content of foods. The analytical method used to quantify may overestimate or underestimate the protein content of the foodstuff. The three most frequently used indirect methods are based on Lowry, Bradford and Kjeldahl techniques [8]. Therefore, the main objective of this study was to document the protein yield, allergenicity and sensitivity in immunoassays of two immunogenic seeds as a result of the choice of the extraction buffer. As the determination of the protein content of the seeds was not the aim of this study only one analytical method [9] was used.

## Materials and Methods

### Peanut protein Extraction

Peanut (Combrasil - 13L02MCN0049) protein extraction was performed as described by Landry [6], and adapted by Teixeira [10]. In short, seeds were milled in an electric coffee grinder (Philco PERFECT COFFEE 127V model 53901040) and passed through a fine stainless-steel sieve. 10g of peanut flour was placed in 4 Falcon tubes. To each of the tubes containing the seed flours we added 100mL qsp (Quantum Satis para) of one of the buffers to be tested. Namely borate buffer (BB), borate buffer with addition of 2% β-mercaptoethanol (Sigma-Aldrich, São Paulo, Brazil, M3701) (BB-2βME), saline buffer (SB) or Tris/HCl buffer (TB/HCl). All 4 tubes were then placed on a rocker at room temperature for an hour. The material of each tube was paper-filtered and the eluted solution (crude extract) was centrifuged three times at 5°C and 2000G for 15 minutes (IEC International Refrigerated Centrifuge Model: PR-2 S/N 26768P-2). At the end of each centrifugation, the top layer (containing fat) and the seed precipitate were discarded. The intermediate supernatants were collected, recentrifuged or aliquoted and stored at −20 ° C until use. Thus, we obtained four crude peanut extracts: crude peanut extract - Borate Buffer (CPE-BB), crude peanut extract - Borate Buffer + 2% β-mercaptoethanol (CPE-BB2βME), crude peanut extract - Saline Buffer (CPE-SB), Crude Peanut Extract - Tris/HCl Buffer (CPE-TB/HCl).

### Protein quantification, SDS-PAGE and immunoblotting

Protein quantification of the crude extracts was determined by the Lowry technique[9]. Protein profile was analyzed using 15% sodium dodecyl sulfate polyacrylamide gel electrophoresis (SDS-PAGE) as described by Laemmli [11] and stained with Coomassie Brilliant Blue (Bio-Rad, Rockville Centre, NY 11571) [12]. Immunoblotting are made to study the immuno-reactivity of the proteins generally as described by Towbin et al.[13]

### In Vivo Experiments

All procedures and number of animals for this work were approved by the university’s ethical committee under the number 781.

Adult (8 weeks old) female C57BL/6 (n=40) and Balb/c (n=40) mice were obtained from the local animal breeding facility (Núcleo de Criação de Animais de Laboratório - NAL – UFF) and kept in the experimental animal-room of the Immunobiology Department during the whole protocol. Animals were kept in polystyrene cages with steel covers in a conventional environment with acidified water and commercial mouse chow *ad libitum* (temperature of 22°C, ~60% humidity and 12-hours light/ 12-hours dark cycle) and were divided in eight groups of C57BL/6 mice and eight groups of Balb/c mice (n=5).

Mice were submitted to a food-allergy induction protocol described by Teixeira (2009). In short 100μg of protein extract was introduced twice subcutaneously with a three-week interval. To the primary inoculation, 1mg of alum (Al(OH)3) was added.

### Immunogenicity/immunoreactivity

Anti-peanut antibodies were tittered using ELISA. In short, each of the 96 wells of the microtiter plates (Global Plast, China, 655111T) was adsorbed with 100μl of saline buffer containing 2μg of the specific protein extract. A 3-fold serial dilution of the sera was performed and HRP goat anti-mouse γ-chain was used (SIGMA, Saint Louis, MO, United States A9044). The result was registered using a microtitration plate reader (Anthos 2010^®^, Biochrom, Cambourne, Cambridge, UK). Reactivity analysis was performed by comparing the area under the dilution curve. To test the cross-reactivity of the different protein extracts the serum of each mouse was tested on microtiter plates adsorbed with one of the 4 protein extracts of the corresponding seed.

### Statistical Analysis

We considered a minimum of five animals per group. The parameters used to display the results were mean and standard deviation. First, Shapiro-Wilk test was used for normal distribution evaluation. Then, outliers were removed using Grubbs test. Finally, the minimum significance level with 95% confidence interval (p) was calculated using Student’s T-test when comparing two groups or one-way analysis of variance (ANOVA) with Tukey-Kramer post-test when comparing multiple groups (multiplicity adjusted p-value). All statistical tests were performed on GraphPad Prism^®^ 6.01 software by GraphPad Software, Inc. (La Jolla, CA, USA).

## RESULTS

### Protein quantification

We observed that the four extraction buffers rendered significantly different protein concentrations for peanuts (Fig 1). The highest CPE yield was obtained with BB2ßME (22,82 ± 1,19 mg/mL) followed by SB (20,94 ± 2,33 mg/mL), BB (18,72 ±2,92 mg/mL) and TB/HCL (12,99 ± 2,32 mg/mL) extraction (Fig 1a). These results show the different extraction potential of each buffer for these two seeds.

**Fig 1.**
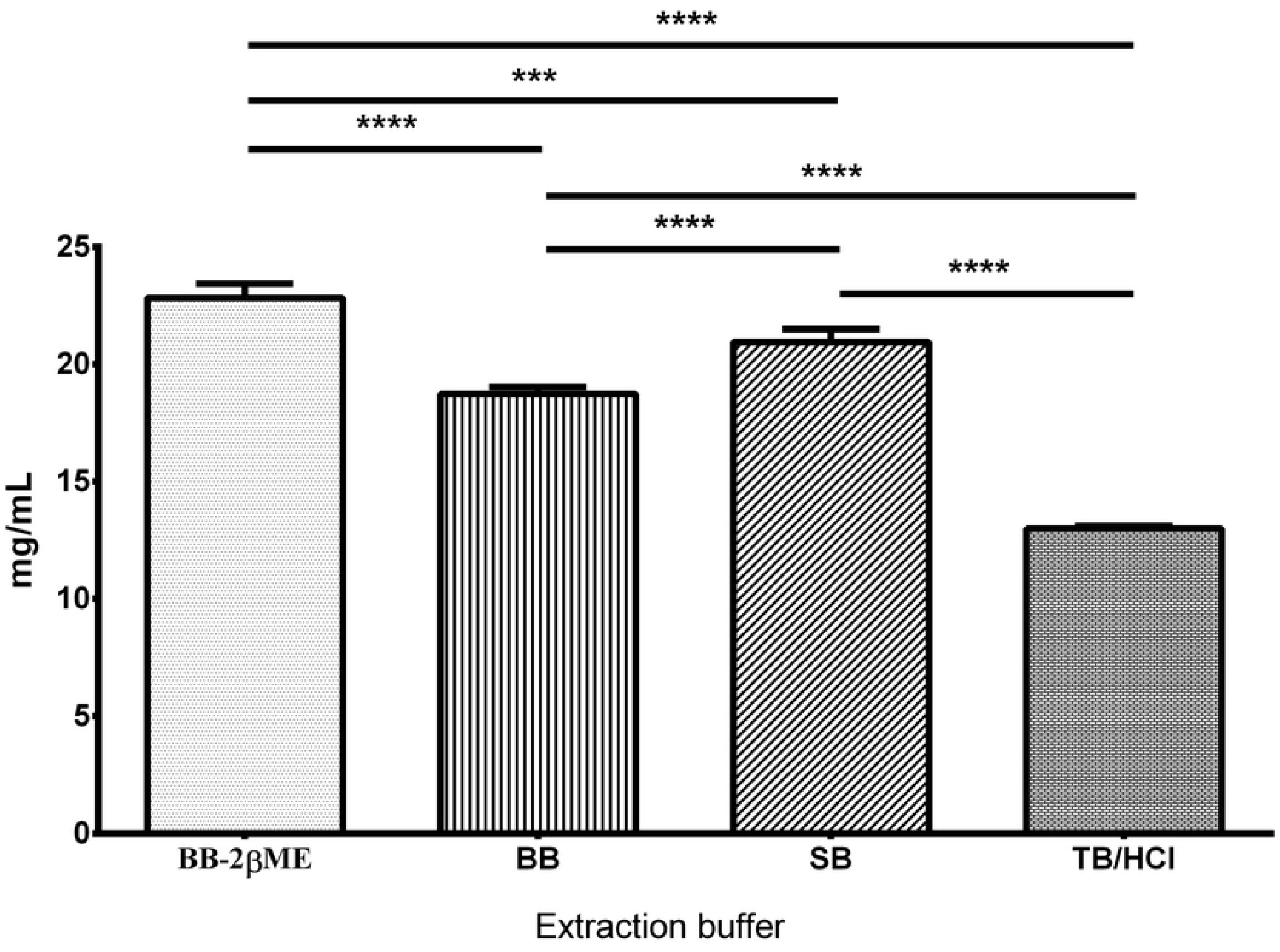
Peanut protein extract yield: Crude peanut protein extracts (CPE) obtained with BB-2βME, BB, SB and TB/HCl; p <0.05 = *; p <0.01 = **; p <0.001 = ***; p <0.0001 = ****

### Electrophoresis - Protein profile determination and immunoblotting

In comparison to BB the use of BB-2ßME led to a shift in the electrophoretic protein pattern of CPE in which some of the higher molecular weight bands reduce in intensity (white double arrow – Fig 2) and others disappear. The pattern of protein bands obtained with SB is very similar to that obtained with BB. However, SB solubilized better the higher and lower molecular weight bands. The immunoblot reactivity showed that the greatest reactivity with sera of animals immunized by CPE-BB-2βME and CPE-TB/HCl to their corresponding extract (Fig. 3).

**Fig 2.**
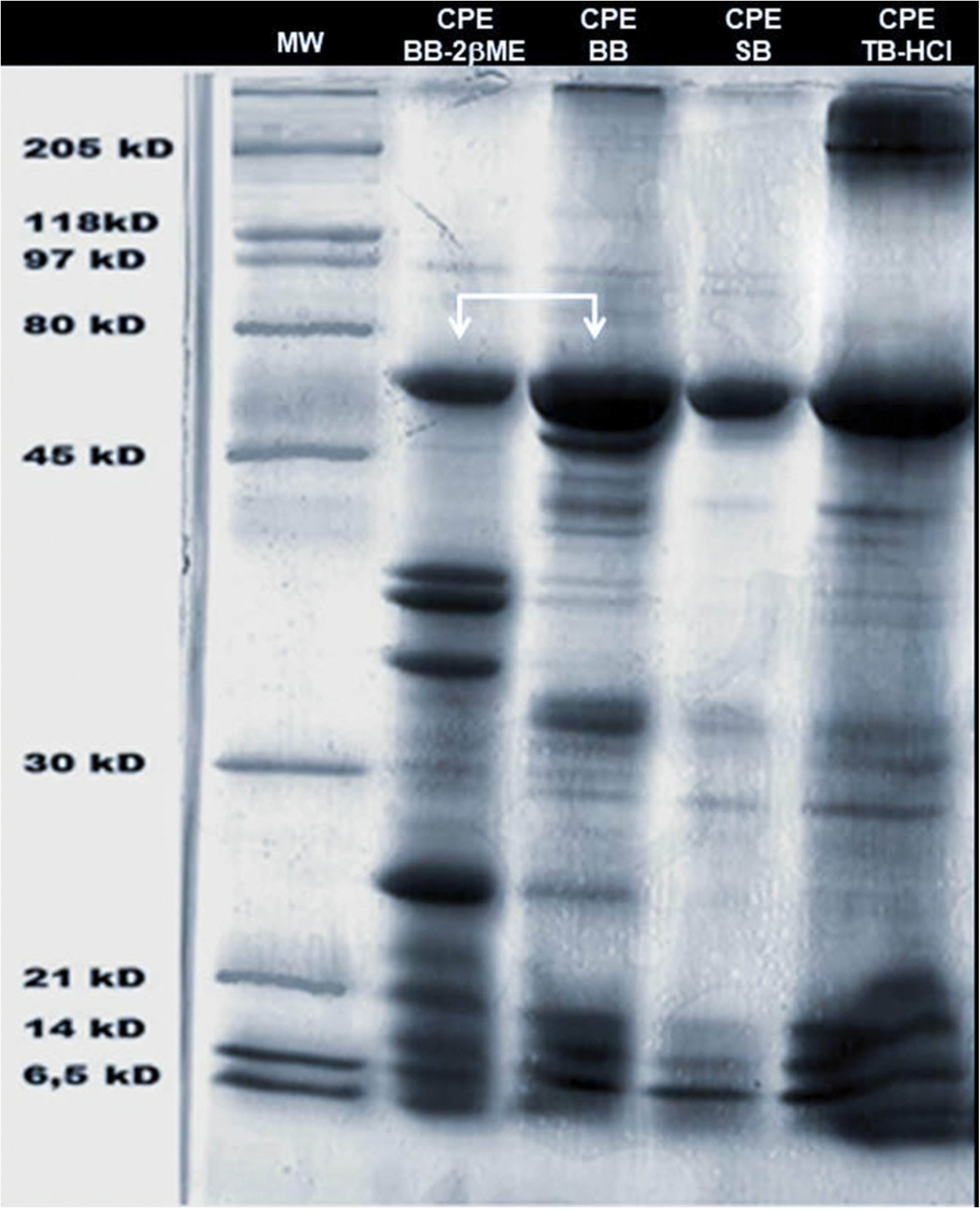
SDS-PAGE - 15% polyacrylamide gels showing the band profile of peanut extracts. The gels were stained with Coomassie Brilliant Blue. The first column shows the molecular weight bands (MW). CPE: Crude peanut protein extracts. BB-2βME: borate buffer with addition of 2% β-mercaptoethanol. BB: borate buffer. SB: Saline buffer. TB/HCl: Tris/HCl buffer.

**Fig 3.**
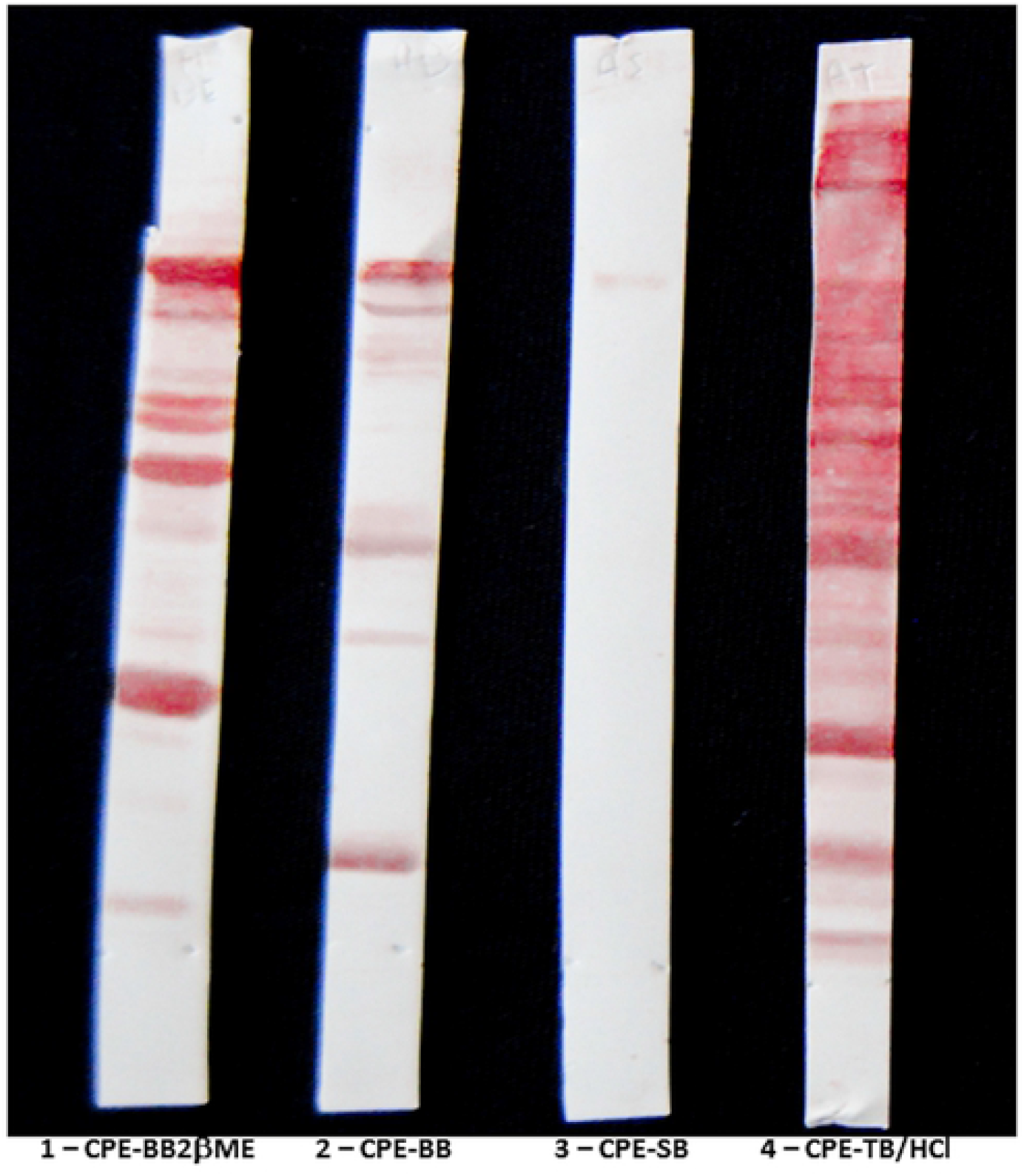
Immunoblotting. Reactivity of a pool of 5 sera to the corresponding sensitization extracts. CPE: Crude peanut protein extracts. BB-2βME: borate buffer with addition of 2% β-mercaptoethanol. BB: borate buffer. SB: Saline buffer. TB/HCl: Tris/HCl buffer.

### Immunogenicity/immunoreactivity

All 4 CPE were immunogenic for both mouse strains tested. All sera showed positive results when reacting to the homologous protein extract immobilized on the ELISA plate. There were no significant intergroup differences in serum reactivity of CPE sensitized C57BL/6 mice (Fig 4A). However significant differences in reactivity of sensitized Balb/c mice were observed (Fig 4B). Animals sensitized with CPE-BB2ßME and those sensitized with CPE-BB presented significantly higher mean antibody titers (p<0.01 and p<0.001, respectively) than those sensitized with CPE-TB/HCl and animals sensitized with CPE-BB presented significantly higher mean antibody titers (p<0.05) than those sensitized with CPE-SB (Fig 4B).

An intriguing observation is the intragroup variation for C57BL/6 mice sensitized with any of the four CPE compared to a lower variation of Balb/c mice to the same protein extracts except for CPE-BB2βME.

**Fig 4.**
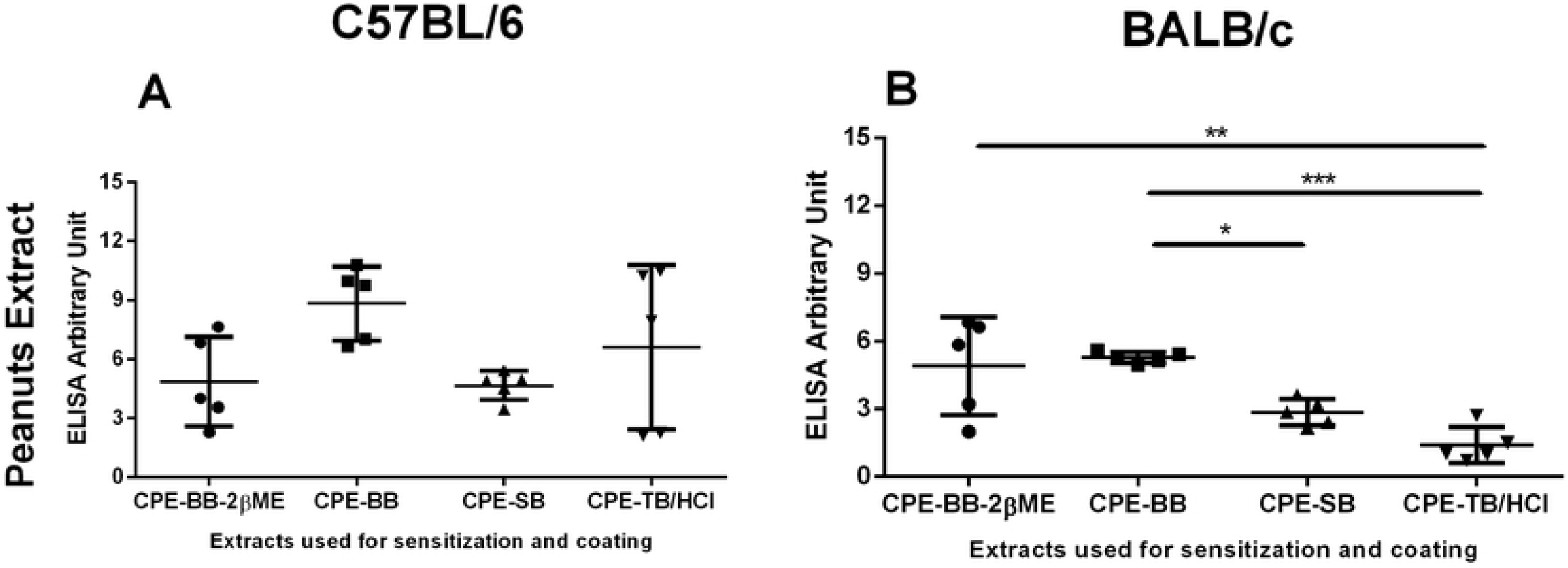
Titer of the sera of C57BL/6 mice (A) and Balb/c mice (B) immunized with Crude peanut extracts. tested on ELISA plates adsorbed with the antigens corresponding to the immunization. N = 5. p <0.05 = *; p <0.01 = **; p <0.001 = ***; p <0.0001 = ****

Next we examined whether different protein extracts of a given seed used to coat polystyrene microtitration plates influenced the antibody titers. Plates coated with CPE-BB2ßME (Fig 5A) or CPE-TB/HCl (Fig 5D) did not distinguish sera reactivity derived from C57BL/6 mice sensitized with any of the four peanut extracts. However, the same sera from CPE-BB2ßME sensitized animals showed significant differences when compared to sera of those sensitized with CPE-BB on plates adsorbed with either CPE-BB (Fig 5B) or CPE-SB (Fig 5C) (p<0.05 and p<0.01, respectively). Thus, the reactivity pattern differs according to the protein extract used to coat the microtitration plates.

**Fig 5.**
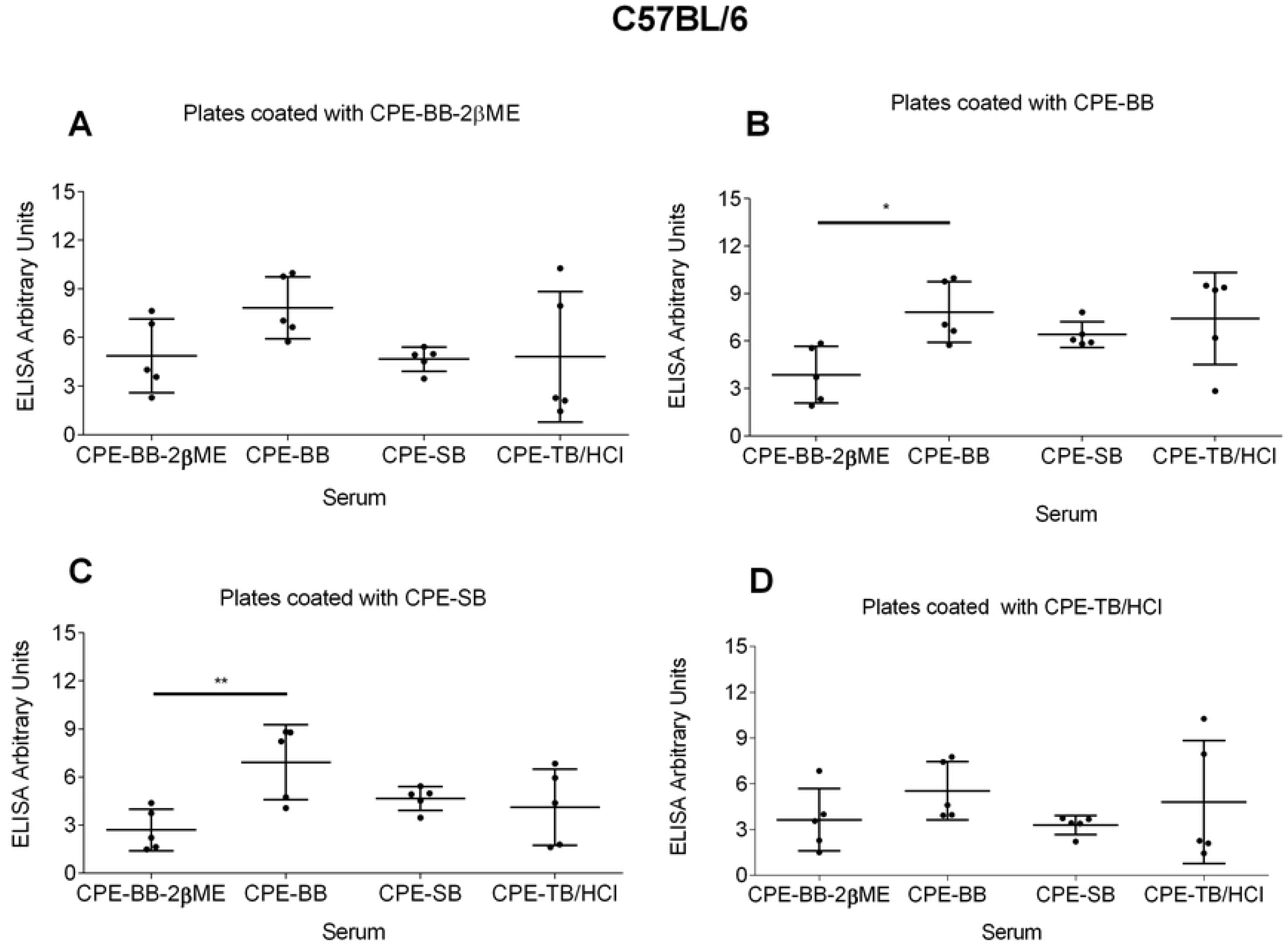
Titers of C57Bl/6 mice sensitized with CPE tested on ELISA. plates adsorbed with A) CPE-BB2βME; B) CPE-BB; C) CPE-SB and D) CPE-TB / HCl. N = 5. p <0.05 = *; p <0.01 = **; p <0.001 = ***; p <0.0001 = ****

Sera from Balb/c mice sensitized with one of the four CPE and tested on CPE-BB2βME adsorbed plates (Fig 6A) showed significant differences in reactivity. Although sera from CPE-BB2ßME and CPE-BB sensitized mice presented similar mean antibody titers the intragroup variation was very different. The first showed a high, while the second a low, titer dispersion. Serum from CPE-BB2ßME and CPE-BB sensitized animals developed significantly higher antibody titers than CPE-SB and CPE-TB/HCl (p<0.001). On CPE-BB adsorbed plates (Fig 6B), serum from CPE-BB developed significantly higher antibody titers than CPE-BB2ßME (p<0.05). The sera from these same four CPE sensitized animals presented no intergroup differences in reactivity when tested on CPE-SB (Fig 6C) or CPE-TB/HCl (Fig 6D) coated plates.

**Fig 6.**
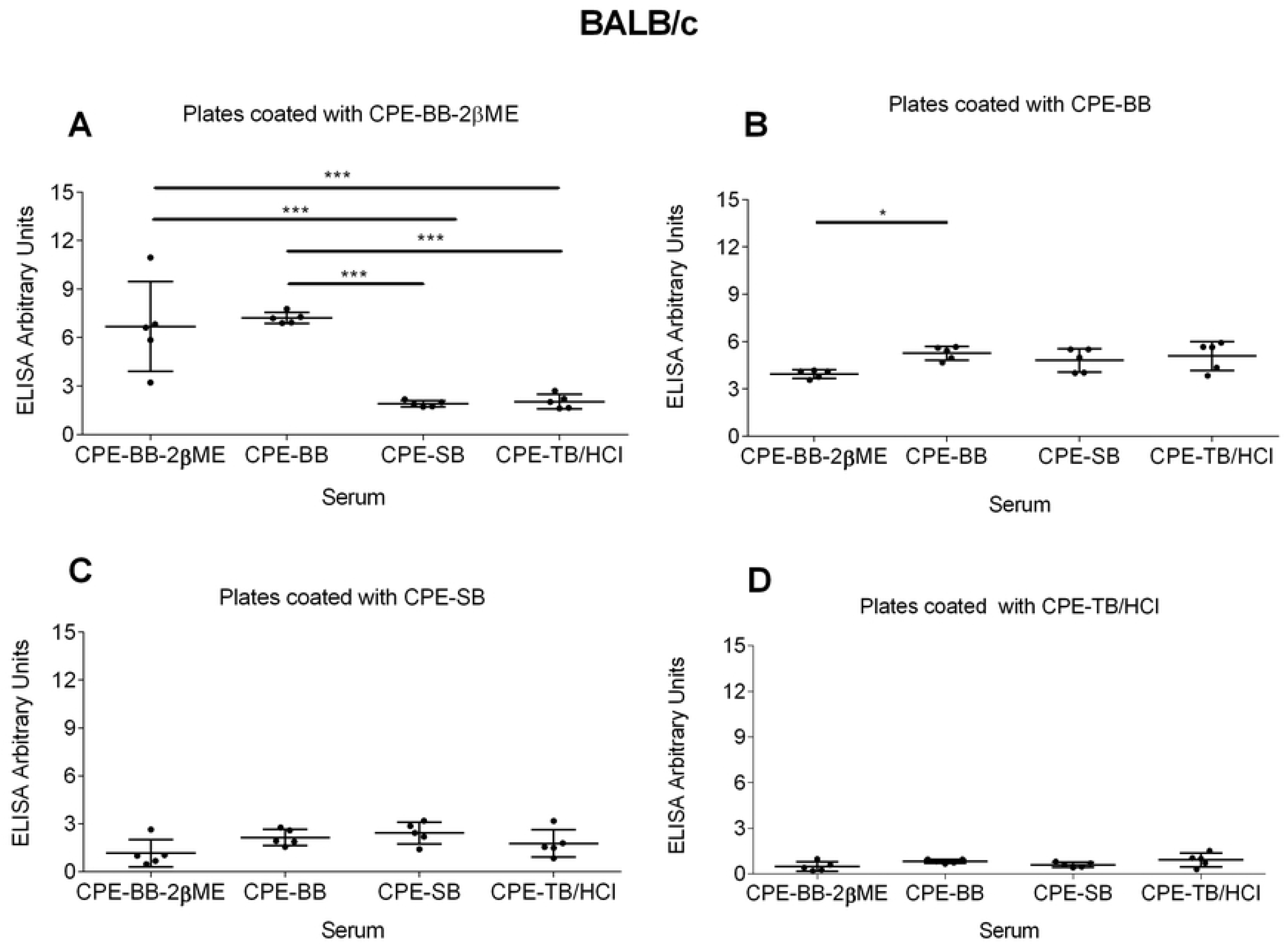
Serum reactivity of Balb/c mice sensitized with peanut extracts tested on ELISA plates. adsorbed with A) CPE-BB2βME; B) CPE-BB; C) CPE-SB and D) CPE-TB / HCl. N = 5. p <0.05 = *; p <0.01 = **; p <0.001 = ***; p <0.0001 = ****

Next, we performed an inter-lineage comparison using the sera derived from Balb/c and C57BL/6 mice that were sensitized and tested on ELISA plates coated with the homologous extract. No inter-strain differences were observed in CPE-BB2ßME, or CPE-TB/HCl reactivity. However, CPE-BB and CPE-SB (p<0.05) induced higher antibody titers in C57BL/6 mice than in Balb/c (Fig 7).

**Fig 7:**
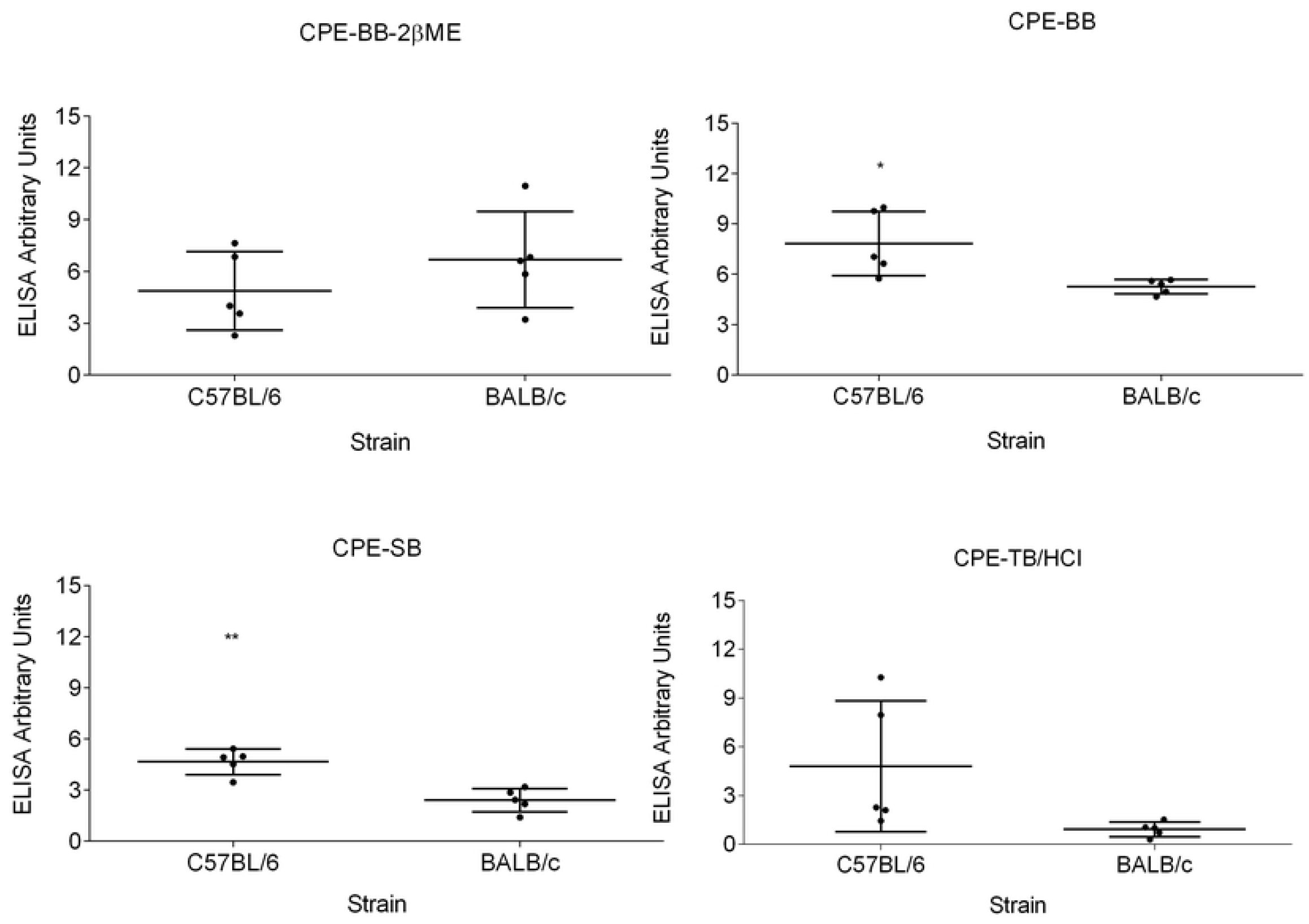
Titers of sera from mice tested on ELISA plates adsorbed with antigens corresponding to immunization. Comparison between C57BL / 6 and Balb/c titers sensitized with CPE. N = 5. p <0.05 = *; p <0.01 = **; p <0.001 = ***; p <0.0001 = ****

Simulating a situation where individuals are sensitized with antigens processed in different ways, and that Elisa Kits preparation is not homogeneous we set up a heat map matrix to observe the reactivity pattern (Fig 8). As discribed in the previous results and sinthesisted here both sensitization protocols and coating of the ELISA plate influence reactivity.

**Fig 8.**
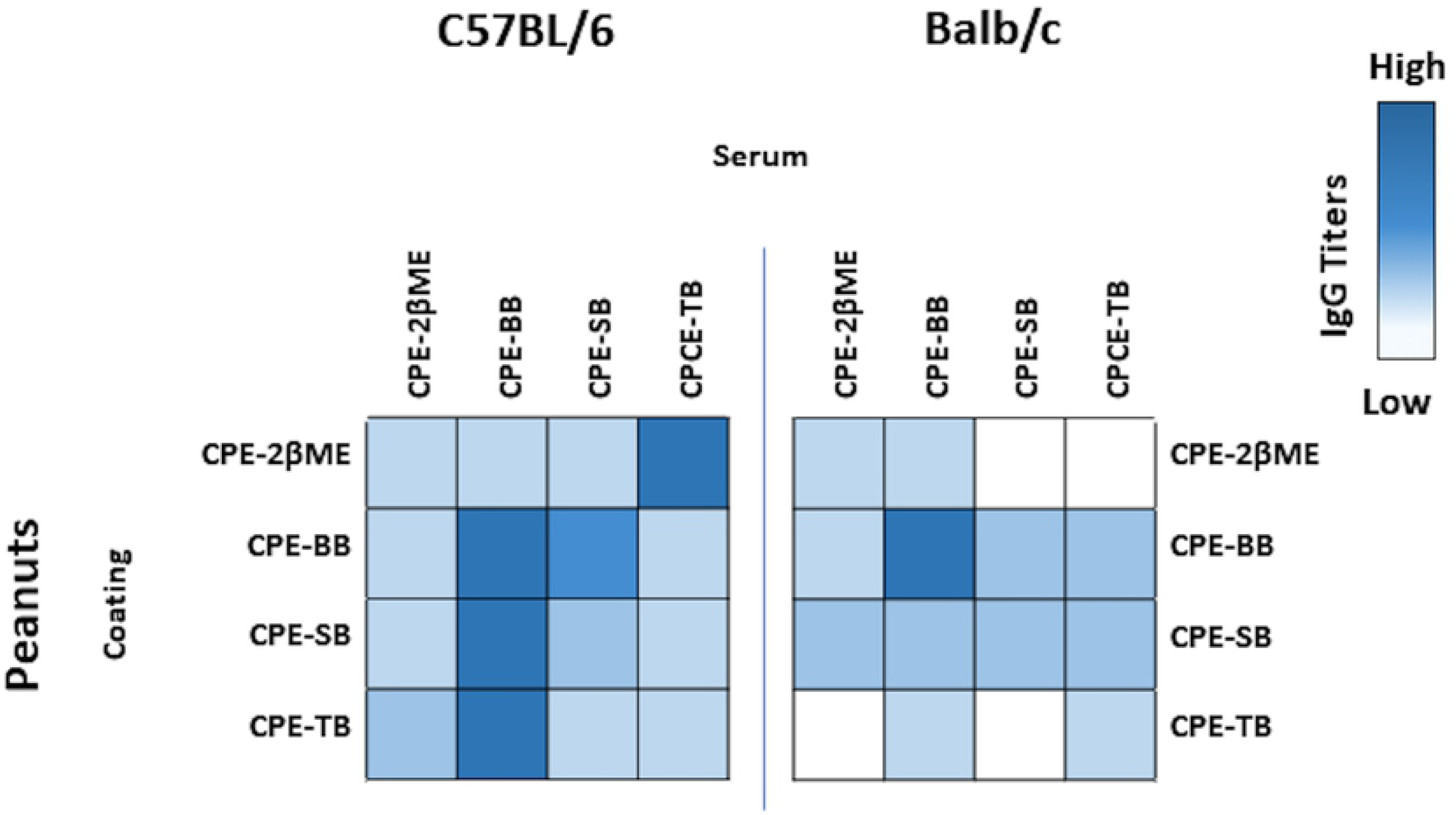
Clustering of anti seed IgG titers. Matrix established crossing administered protein, protein used to coat ELISA plates and mouse strain.

## DISCUSSION

A diet containing immunogenic proteins is extremely important for the maintenance of the homeostasis of the immune system considering the following facts: 1) the maintenance of lymphoid cells in the periphery demands their activation or stimulation through the interaction of their antigen receptor (TCR - T cell receptor BCR – B cell receptor); 2) the TCR interacts with peptides present on the MHC molecules of antigen presenting cells (APC); 3) that most of the lymphocytes of the organism are located in the mucosa and 4) that mucosal lymphoid tissue (MALT) is the largest interface of the immune system with the external environment. A formal demonstration of this latter fact showed that animals fed on an elemental (monomeric) diet as of weaning, therefore with no immunological stimulus, presented, in adult life, a significant decrease of their MALT, characterized by the reduction in the number of intraepithelial lymphocytes, cells of the lamina propria and Peyer patches. The size and cell count in other lymphoid organs was also lower [14]. But what makes a dietary protein a food allergen? This has not yet been fully established. The knowledge about the factors that determine the allergenicity of food proteins is still incomplete. In theory any protein contained in foods could induce allergy, however, as stated before 14 are responsible for over 90% of all allergies.

From the antigenicity point of view peanut is one of the best studied legumes. The World Health Organization recognizes 17 distinct peanut allergens.

In order to perform *in vivo* or *in vitro* diagnostic tests purified antigens are fundamental. Early characterization of peanut allergen utilized serum from allergic patients to prepare affinity columns. The study of the eluted proteins permitted the molecular characterization of the major allergens [15]. Since the 1980’s several molecular biology strategies were developed and are currently being used to produce purified proteins both from food *in natura* as well as recombinant proteins. Albeit all efforts there are still inconsistencies between manufacturers of food allergens which may turn the tests unreliable in the clinical setting [16].

We prepared peanut extracts using four different buffers ranging in pH from 6.8 to 10.0. Quantitively, the highest yield was obtained with the alkaline borate buffer with the addition of β-mercaptoethanol, a disulfide bond reducing agent. Regarding the other three buffers we observed that the use of a borate buffer without the addition of β-mercaptoethanol extracted fewer peanut proteins than the neutral pH saline buffer. Thus, the neutral SB extracted more peanut protein than both alkaline BB and acid TB. We did not find in the literature explanation for this fact.

The Brazilian Table of Food Composition (TACO) research group obtained each food from the five Brazilian macro regions, then analyzed its micro and macro elements [17]. The food table refers to the average value of each component contained in the respective foods. Protein quantification was performed by the Kjeldahl method that determines the total nitrogen contents which was then converted to protein concentration [18–20]. Our results using the Lowry [9] protein quantification method agree with the TACO results [17]. We can argue that the use of β-mercaptoethanol may have better exposed reactive sites of the protein content than the other three buffers.

Having obtained the protein extracts we determined the allergenicity of each extract by performing a routine sensitization protocol with alum, an adjuvant typically used to induce IgE [21] in contrast to Freud adjuvant that is typically used to stimulate IgG [22]. The two mouse strains used - C57BL/6 and Balb/c - were chosen due to their Th1 and Th2 profiles, respectively. To study the intrinsic immunogenicity of allergens and the influence of the food matrix, van Wijk, Nierkens (23) compared the *in vivo* (subcutaneous) response to purified peanut allergens and to crud peanut extract (PE) similar to our extract. These authors inform that none of the purified allergens induced significant immune activation while, the extract induced an increased immune response to the individual allergens. In accordance to the authors, one must argue that purified proteins injected in the subcutaneous tissue without an inducer of inflammation may be immunogenic in the long run (for example the use of porcine insulin by diabetic patients) but most probably not after a single inoculation.

Although defatting the milled seeds is a common procedure in protein purification, we did not include this step prior to the extraction procedures, taking in account that the ingested food is, in general, not in a defatted condition. However, during the incubation period and centrifugation process most of the fat was removed as it comes out of phase. The amount of reminiscent fat may have influenced the antigenicity of each extract. A frequently used immunopotentiator is incomplete Freund’s adjuvant which is an antigen solution emulsified in mineral oil [24]. The possible influence of the different buffers on the separation of the fat contained in the seeds was not calculated.

Only to site a few of the potential influencers of allergenicity are: the food matrix [23], protein stability during processing [25], and the immunological status of the entry port [26]. Food matrix can be defined as the way the chemical components of food (protein, fatty acids, carbohydrates, and micronutrients) interact to establish the structural organization from microscopic to macroscopic scales. Both cooked and raw food undergo harsh physical and chemical reactions before it can be absorbed by the gut. The majority must be broken down to the elemental components for absorption, however, to induce allergy some of the molecules must resist processing to continue immunogenic. The immunological status of the gut, as has been previously demonstrated by our group, is essential to induce either allergy or tolerance after the initial contact of the food with the mucosa. [26]

Our data agree with those of the literature [27, 28] regarding the pH of the buffer used in the extraction of peanut proteins - neutral pH is more efficient. Different from these authors the use of the reducing agent increased the extraction efficiency of our model. The difference between the two protocols consists in the buffer and the reducing agents used. We used an acid while they used an alkaline tris buffer and we used βME and they used SDS and other reducing agents. Our data are in accordance to the literature. As proposed by these authors the introduction of crude peanut extract is immunogenic. Crude peanut extract consists of part of the proteins, carbohydrates and fatty acids derived from the seed which may be responsible for the adjuvant effect due to their inflammatory potential. [23].

In our experiment, all protein extracts are immunogenic. All animals (a total of 40 = 4 extracts, 2 mouse strains, 5 animals per group) produced significantly more antibodies to the inoculated protein-adjuvant (alum) solution, than their sham counterparts. Irrespective of which protein extract was inoculated or adsorbed to the Elisa plate positive reactivity was observed. However, the intensity of serum reactivity varied. C57BL/6 mice sensitized with one of the four peanut extracts did not show significant differences when the plates were adsorbed with CPE-BB2ßME or CPE-TB/HCl. These results suggest a dominance in the allergenicity profile of the bands that were obtained by these two buffers. On the other hand, the same sera show distinct reactivity profiles, as can be seen in our results, when the plates are coated with CPE-BB or CPE-SB.

As can be seen in the electrophoresis the profile of protein bands is influenced by the extraction buffer used. Although similar protein bands are extracted by each of the extraction buffers there are bands that are not present in all extracts. Two hypotheses arise, either the remaining bands (those that are unique for each extract) are not as immunogenic or the proteins that adsorbed to the titration plates derived with these buffers have similar profiles. These hypotheses were not tested. We are currently undertaking further experiments to elucidate this fact.

Each crude protein extract induced different intragroup variability. C57BL/6 mice sensitized with CPE-SB presented more homogeneous reactivities when compared to those sensitized with CPE-TB/HCl which presented quite heterogeneous reactivities. Some individuals with high levels and others with low levels of anti-peanut antibodies. This pattern of response was less prevalent in Balb/c animals, being visualized only in animals inoculated with CPE-2βME and tested with plates adsorbed with the homologous extract.

In populations with a heterogeneous genetic profile one would expect a pattern of intragroup discrepancies however the experiments were undertaken with inbred mouse strains. We found no similar findings in the literature and our working hypothesis is that the animals’ microbiota may be influencing their responses. In the room that houses the experimental animals in our animal facility the cages are intentionally maintained on open shelves, thus subject to environmental influences as in “real life”. We have preliminary results from other experiments supporting our findings, however they still do not adequately explain our data.

The mean reactivity of sera from the C57BL/6 and Balb/c mice sensitized with one of the four peanut extracts varied according to the extract used for adsorption on the plates. The mean reactivity titers of sera plated on CPE-2βME crude protein extracts were lower than the remaining plates suggesting that either its adhesion is less effective or that 2βME decreases the antigenic capacity of this extract as it dissociates disulfide bonds. The latter is supported by the fact that immunizations with this extract also yields lower titers on all other plates covered with the extracts obtained with other buffers (CPE-BB, CPE-SB and CPE-TB / HCl).

The immunoblotting revealed that the pattern of reactivity of the sera to the corresponding extract used to sensitize also varies according to the crude extract used. Astuti et al in their work found that peanut allergen induced individual specificity patterens, suggesting that it is better to use crude protein extracts rather than purified proteins as a tool for diagnosing allergies, since immunodominance may vary due to food processing, absorption and physiology therefore, each individual may be allergic to different allergens of any food [29].

We did not observe differences in the comparison of the mean reactivities of sera from C57BL/6 mice sensitized with one of the four peanut extracts and tested on an ELISA plate adsorbed with the homologous extract. In this lineage although the average reactivity for the peanut was not significantly affected it is possible to observe the great intragroup dispersion with some animals responding more than others. This observation is not the result of possible inter-plate variations because all the sera from a same group were evaluated on the same plate suggesting the interference of other biological factors in the response to the peanut by C57BL/6 mice.

This scenario is different for the Balb/c mice sensitized with the peanut extracts. The form of extraction clearly alters the reactivity of the CPE-TB/HCl showed the lowest titers. In this lineage only those inoculated with CPE-BB2βME showed a large dispersion in anti-peanut IgG titers. Although CPE-TB/HCl yielded the lowest protein concentration, this fact does not justify the low immunization since all the animals were inoculated with 100 μg of antigen independent of the extract, suggesting that genetic factors of the lineage associated with the differentiated immunogenicity of the extracts influenced the observed immune response.

Among the most commonly used mouse strains in immunological research, are the C57BL/6 and Balb/c mice. These typically present a Th1 and Th2 profile respectively. Stanisavljević [22] demonstrated that intradermal immunization of mice with complete Freund’s adjuvant resulted in alteration of the distribution of Th cells in the GALT of C57BL/6 mice, but not of Balb/c mice that responded poorly to immunization [30].Demonstrated different responses between C57BL/6 and Balb/c mice submitted to an epicutaneous exposure of antigens that sensitizes for experimental eosinophilic esophagitis in the former but not the latter.

In order to verify some association with the genetic factors, a comparison of the different sensitizations between the C57BL/6 and Balb/c lines was carried out. As was demonstrated the reactivity profile differs due to both strain and foodstuff.

## Conclusion

The addition of β-mercaptoethanol to the borate buffer ensures a greater efficiency in extracting the proteins of peanut and the Tris/HCl buffer does not have a great efficiency in the extraction of proteins from peanuts. The protein profile varies according to the extraction buffer used and all four buffers were suitable for the extraction of immunogenic proteins from peanuts. The use of different protein profiles can prevent false negatives. On the other hand, its unrestricted use can lead to false positives.

## Acknowledgments

The authors would like to thank the Coordenação de Aperfeiçoamento de Pessoal de Nível Superior (CAPES) - Coordination for the Improvement of Higher Education Personnel (CAPES) and the Conselho Nacional de Desenvolvimento Científico e Tecnológico (CNPq) - National Council for Scientific and Technological Development (CNPq) for their financial support.

